# Bacteriophages targeting protective commensals impair resistance against *Salmonella* Typhimurium infection in gnotobiotic mice

**DOI:** 10.1101/2022.09.28.509654

**Authors:** Alexandra von Strempel, Anna S. Weiss, Johannes Wittmann, Marta Salvado Silva, Diana Ring, Esther Wortmann, Thomas Clavel, Laurent Debarbieux, Karin Kleigrewe, Bärbel Stecher

## Abstract

Gut microbial communities protect the host against a variety of major human gastrointestinal pathogens. Bacteriophages (phages) are ubiquitous in nature and frequently ingested via food and drinking water. Moreover, they are an attractive tool for microbiome engineering due to the lack of known serious adverse effects on the host. However, the functional role of phages within the gastrointestinal microbiome remain poorly understood. Here, we investigated the effects of microbiota-directed phages on infection with the human enteric pathogen *Salmonella enterica* serovar Typhimurium (*S*. Tm), using a gnotobiotic mouse model (OMM^12^) for colonization resistance (CR). We show that phage cocktails targeting *Escherichia coli* and *Enterococcus faecalis* acted in a strain-specific manner. They transiently reduced the population density of their respective target before establishing coexistence for up to 9 days. Infection susceptibility to *S*. Tm was markedly increased at an early time point after phage challenge. Surprisingly, OMM^12^ mice were more susceptible 7 days after a single phage inoculation, when the targeted bacterial populations were back to pre-phage administration density. The presence of phages that dynamically modulates the density of protective members of the gut microbiota provides opportunities for invasion of bacterial pathogens.

## Introduction

The human intestine is a complex and highly dynamic microbial ecosystem, which consists of trillions of microorganisms, predominantly bacteria (1). In addition to other important functions, the gut microbiome forms a protective barrier against human pathogens such as *Salmonella*, *Clostridioides difficile* or multi-resistant Gram-negative Bacteria (MRGN), termed colonization resistance (CR). Mechanisms underlying CR include substrate competition, production of bacteriocins or toxic metabolites or initiation of host immune responses (2). CR is higher in hosts with a complex microbiota and generally disrupted by drugs (3), dietary factors (4, 5) and diseases (6), but was also shown to be highly variable among healthy human individuals (7, 8). Other risk factors may exist, which are currently still unknown.

Besides bacteria, the human microbiome also includes viruses, amongst which bacteriophages, the viruses that predate on bacteria, are highly abundant (9, 10). Previous studies reported that phages can impact the bacterial community (11, 12) and influence bacterial population structures, functions and the metabolome by interacting with their bacterial hosts. Metagenomic studies identified largely temperate phages in the gut microbiota (13) and showed that their activation can be triggered by various environmental inducers including diet (14, 15). Virulent phages are also found in the mammalian gut (16) where they can coexist with their host bacteria over time (17) and thereby also functionally impact bacterial communities (11). Although metagenomic studies have addressed the abundance, diversity, individuality and stability of phages in the gut (18), little is known about the role of virulent phages in regulating intestinal microbiome functions.

In this work, we investigated the effect of two virulent phage cocktails targeting *Escherichia coli* and *Enterococcus faecalis* on the microbial community and CR against human pathogenic *Salmonella enterica* serovar Typhimurium (*S*. Tm). We used a gnotobiotic mouse model for CR, based on the Oligo-Mouse-Microbiota (OMM^12^), a synthetic bacterial community for functional microbiome research (19). The OMM^12^ consists of 12 bacterial strains representing the five major phyla in the mouse gut, which was recently characterized *in vitro* (20) and can be extended by bacterial strains to provide additive functions. Here, we amended the OMM^12^ with two additional members, *E. coli* Mt1B1 and the secondary bile acid producer *Extibacter muris* DSM 28560 to enhance CR against gastrointestinal pathogens (OMM^14^) (21). Recent studies showed that *E. coli* Mt1B1 and *E. faecalis* KB1 are involved in mediating CR against *S*. Tm in the community context. These two bacterial species are frequently challenged by virulent phages present in the environment and that could reach the gut via drinking water and food (22, 23). We addressed the impact of phages targeting *E. coli* and *E. faecalis* on microbiome functions *in vivo* and *in vitro* and their global effect on the members of a synthetic gut community. By targeting bacteria that play a role in mediating CR against *S*. Tm, we show that phages impair CR and facilitate pathogen invasion.

## Results

### Phage isolation and characterization

To target *E. coli* Mt1B1 we selected three virulent phages, Mt1B1_P3, Mt1B1_P10 and Mt1B1_P17 (short: P3, P10 and P17), which were previously characterized and shown to stably coexist with their host strain in gnotobiotic mice (17). P3 and P10 are podoviruses and belong to the genus *Teseptimavirus* and *Zindervirus*, respectively, whereas P17 shows characteristic features of a myovirus (**Table 1**). In accordance with Lourenco et al. (17), we found that all three phages individually inhibited *E. coli* Mt1B1 growth *in vitro* and showed strongest inhibition when added together (**S1A Fig**). In previous studies, the host range of the individual phages was tested against several different *E. coli* strains (17, 24). It was shown that phage P17 exhibited a rather broad host range whereas the host range of P3 and P10 was narrow.

**Table 1.**
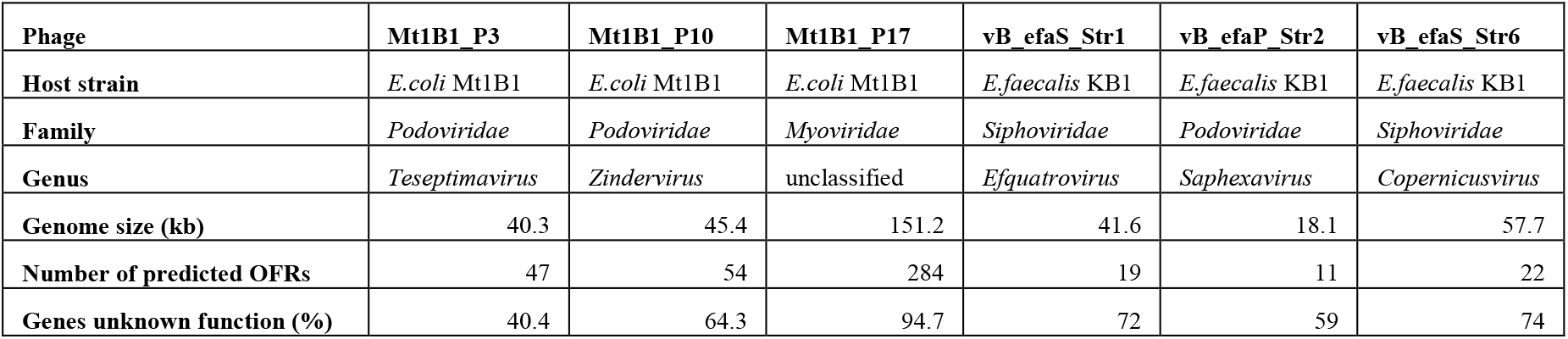

Further, we isolated three phages targeting *E. faecalis* KB1 (DSM 32036) from sewage water: vB_EfaS_Strempel1 (short: Str1, DSM 110103), vB_EfaP_Strempel2 (Str2, DSM 110104), vB_EfaS_Strempel6 (Str6, DSM 110108) and characterized them taxonomically and functionally. Str1 and Str6 are members of the genus *Efquatrovirus* and *Saphexavirus*, respectively, and show characteristic features of siphoviruses, whilst Str2 is a podovirus and belongs to the *Copernicusvirus* genus (**Table 1**). In liquid culture, the phages inhibited *E. faecalis* KB1 growth with different lysis profiles. All three phages added together mostly resemble the lysis behavior of Str1 (**S1B Fig**). Str1 was highly strain-specific and only produced visible plaques on *E. faecalis* KB1 when tested against 12 other *Enterococcus faecalis* and *Enterococcus faecium* strains (**S1C Fig**). In contrast, Str6 exhibited the broadest host range, targeting 5 out of 12 isolates, whereas Str2 lysed 3 out of 12 (**S1C Fig**). In addition, none of the six phages infect the *Salmonella* strain used the following experiments.

### Phages specifically target their host strains within an in vitro bacterial community

We next explored the effect of two phage cocktails (3Φ^Mt1B1^ and 3Φ^KB1^) on a synthetic community harboring *E. coli* Mt1B1 and *E. faecalis* KB1 *in vitro*. To this end, we added *E. coli* Mt1B1 and *E. muris* JM-40 (DSM 28560) to the OMM^12^ community (19, 25) **(S2A Fig**). The latter is a secondary bile acid producer, which is a gut microbiota function lacking in the original OMM^12^ community. The 14 community members (OMM^14^) were added at equal ratios (OD_600_) to anaerobic media (AF medium, (20)) and diluted (1:100) every 24 h in batch culture. On day three, 12 h after dilution, phages were introduced into the batch culture with a rough multiplicity of infection (MOI) of 0.01 and incubation was prolonged for three days with dilution every 24 h (**Fig 1A**).

**Fig. 1:**
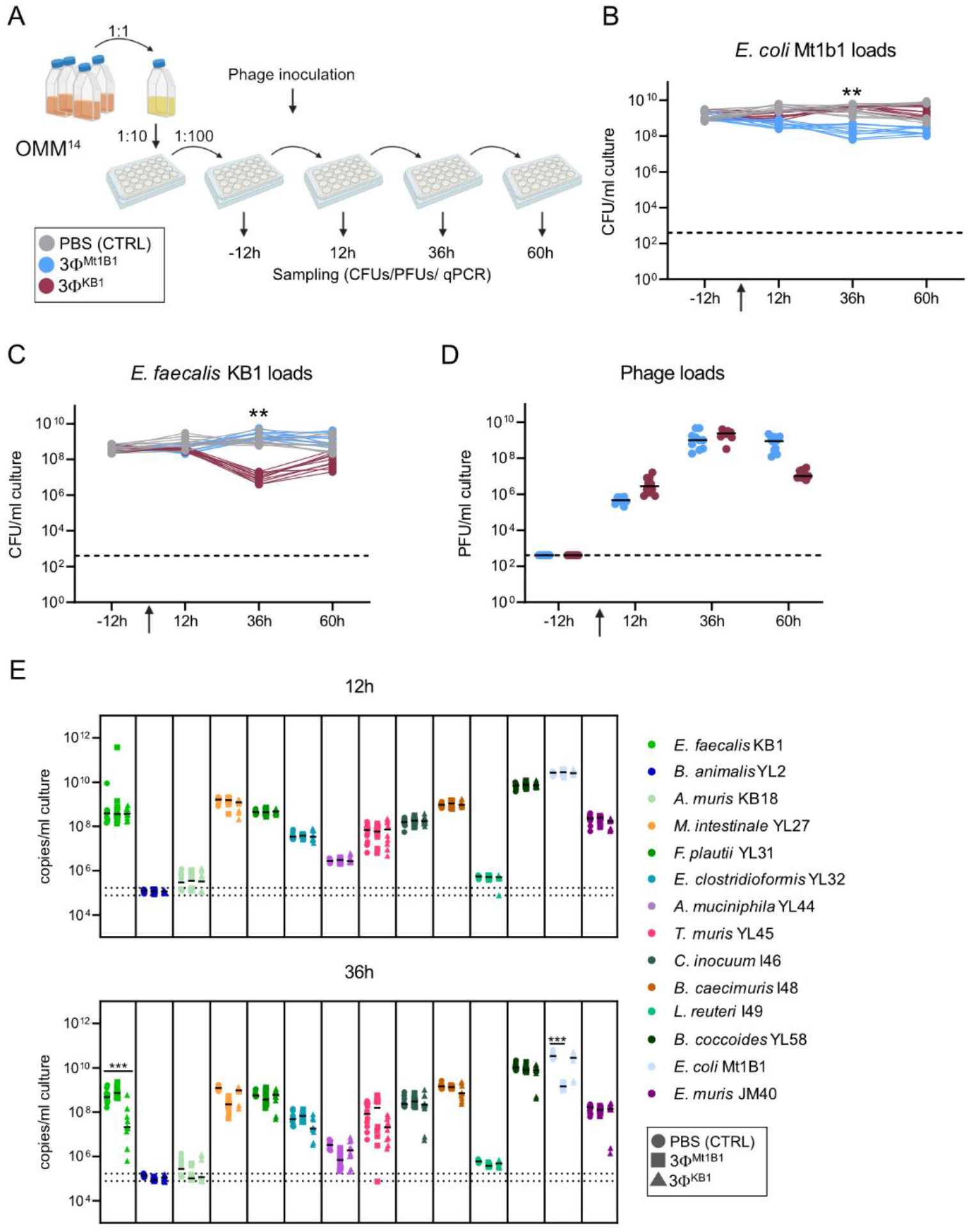
Phages specifically target their hosts in vivo. **(A)** Experimental setup batch culture. The OMM^14^ strains were grown in monoculture, mixed at the same OD_600_ ratio and diluted every day 1:100 in AF medium. 12 hours after the third passage, phage cocktails 3Φ^Mt1B1^ or 3Φ^KB1^ were added to the wells of a 24 well microtiter plate. Before each dilution, samples were taken for plating and qPCR. **(B)** *E. coli* Mt1B1 and **(C)** *E. faecalis* KB1 loads determined by plating (Log_10_ CFU/ml). **(D)** Phage loads were determined via spot assays on host strains *E. coli* Mt1B1 or *E. faecalis* KB1, respectively. **(E)** Community composition 12 h and 36 h after phage addition, absolute abundance of each strain was determined using a strain-specific qPCR and plotted as 16S rRNA copy numbers per ml culture. Statistical analysis was performed using the Mann-Whitney Test comparing the treatment groups (N=10) against the control group (N=10) (* p<0.05, ** p<0.01, *** p<0.001). Each dot represents one well, black lines indicate median, dotted lines indicate detection limit (DTL). The experiment was conducted in two biological replicates with ten technical replicates in total.

A significant drop in *E. coli* CFUs was observed 36h after phage cocktails treatment, followed by an increase of *E. coli* density after 60 h (**Fig 1B**). *E. faecalis* levels in cultures treated with phages also dropped significantly by approximately two orders of magnitude below control levels after 36 h, also resulting in a regrowth after 60 h (**Fig 1C**). All phage titers increased after 36 h, coinciding with the drop in their host bacterial populations (**Fig 1D**). Furthermore, phages were also tracked by specific qPCR, revealing that all three *E. coli* phages as well as *E. faecalis* phages Str1 and Str2 replicate in the batch culture context (**S1D Fig**), whereas phage Str6 becomes undetectable after 60 h (**S1E Fig**).

The community composition was monitored by qPCR and remained stable over six days and between the replicates (**Fig 1E, S1F Fig**). The species *E. coli* and *Blautia coccoides* YL58 dominated the communities in absolute abundances as 16S rRNA copies per ml culture, as previously observed (19-21, 26). All strains but *Bifidobacterium animalis* YL2, *Acutalibacter muris* KB18 and *Limosilactobacillus reuteri* I49 were detected and showed minimal changes in their absolute abundance before and after phage treatment (**Fig 1E**). The overall stability of the OMM^12^ community members was not affected by the addition of either of the two phage cocktails (**Fig 1E, S1F Fig**) suggesting that these phages can be used as strain-specific tools for community manipulation.

### *E. coli* Mt1B1 and *E. faecalis* KB1 are specifically targeted by phage-cocktails in gnotobiotic OMM^14^ mice

Next, we investigated effects of the phage cocktails (3Φ^Mt1B1^ and 3Φ^KB1^) on their host bacteria *in vivo*. To this end, we established a stable colony of gnotobiotic OMM^14^ mice. Twelve out of the 14 bacteria colonized the mouse gut in a stable manner over several generations, as quantified in feces by qPCR (**S2B Fig**). *B. longum* subsp. *animalis* YL2 and *A. muris* KB18 were not detected, as previously shown in OMM^12^ mice (24, 26); these strains either do not colonize or fecal levels are below the detection limit of the qPCR.

Using the OMM^14^ mouse model, we studied the microbial community composition in response to a single oral challenge of 3Φ^Mt1B1^ or 3Φ^KB1^ (1 × 10^7^ PFU of each phage in 100μl PBS or only PBS as control without phages) in OMM^14^ mice (n = 4-6, **Fig 2A**). Absolute abundances of the targeted bacteria and their phages in the feces was monitored on day 1 throughout 4 and day 7 post phage challenge (p.c.) via strain- and phage-specific qPCR. The treatment with 3Φ^Mt1B1^ significantly reduced *E. coli* Mt1B1 loads on day 1-4, but strain 16S rRNA gene copy numbers had reached again basal levels by day 7 (**Fig 2C**). The reduction of *E. faecalis* KB1 loads was more pronounced at day 2 p.c. in mice treated with 3Φ^KB1^, but the population also recovered by day 7 p.c. (**Fig 2D**). The absolute abundances of the 12 other OMM members were not significantly affected by phage administration (**Fig 2E; S3A Fig**) and remains more stable compared to the batch culture.

**Fig. 2:**
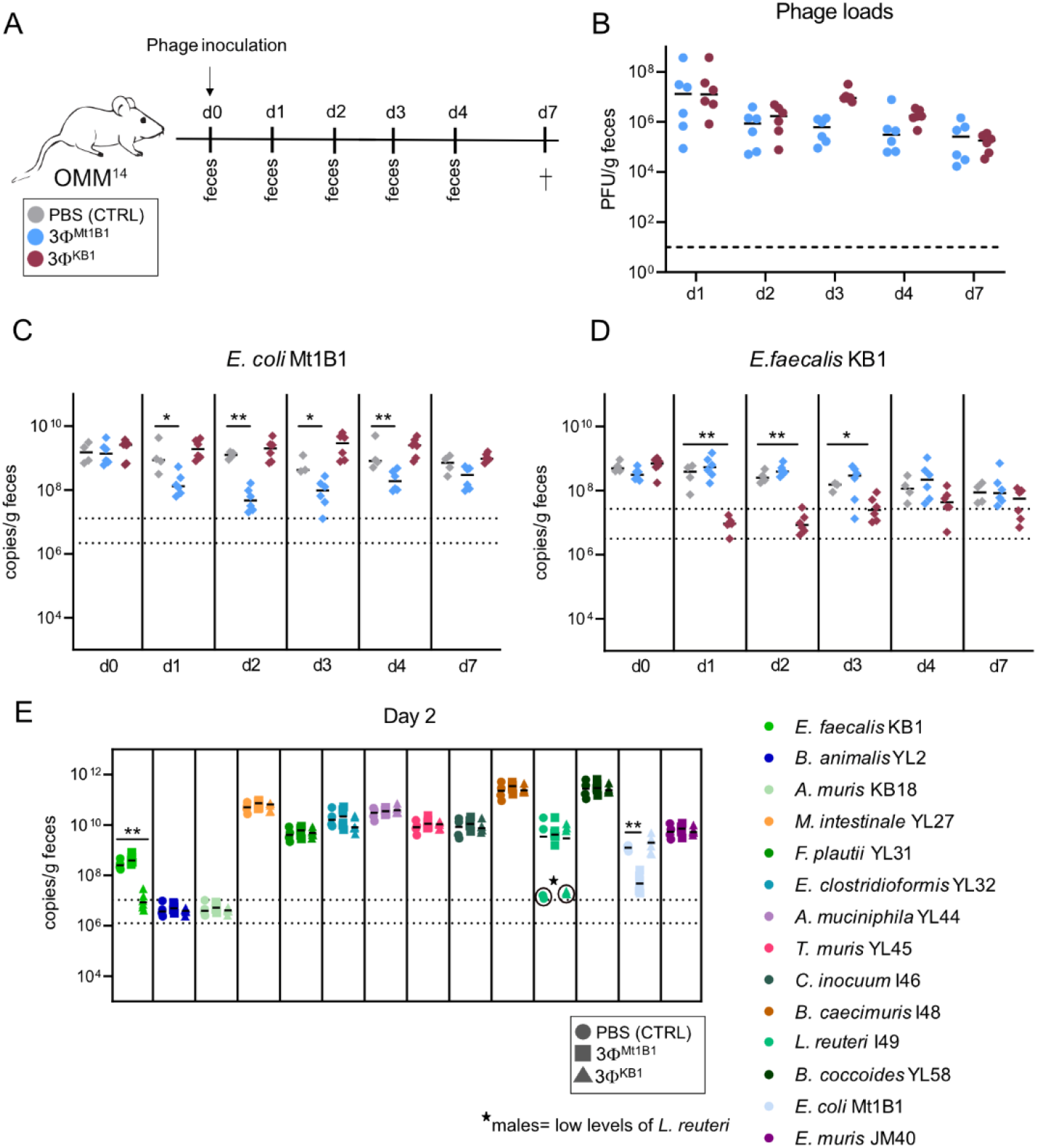
*E. coli* Mt1B1 and *E. faecalis* KB1 can be specifically targeted by phage-cocktails *in vivo* in OMM^14^ mice. **(A)** Experimental setup, mice stably colonized with the OMM^14^ community were challenged orally with phage cocktails 3Φ^Mt1B1^ or 3Φ^KB1^ or PBS as control (10^7^ PFU per phage) and feces were collected every day. On day 7 post phage challenge (p.c.), the mice were sacrificed. **(B)** Phage loads (PFU/g feces) were determined via spot assays. **(C)** *E. coli* Mt1B1 and **(D)** *E*.*faecalis* KB1 loads were determined by qPCR in fecal samples at different time points p.c.. **(E)** Absolute quantification of all other members of the OMM^14^ consortium on day 2 p.c. by strain-specific qPCR (16S rRNA copies/g feces). Statistical analysis was performed using the Mann-Whitney Test comparing the treatment groups (N=6) against the control group (N=4) (* p<0.05, ** p<0.01, *** p<0.001). Each dot represents one mouse, black lines indicate median, dotted lines indicate limit of detection.

Total phage levels determined as PFU per g feces were comparable for 3Φ^Mt1B1^ and 3Φ^KB1^, reaching a maximum on day 1 p.c. with approximately 10^7^ PFU/g feces, followed by stable levels between 10^5^ PFU/g feces −10^6^ PFU/g feces until day 7 p.c. (**Fig 2B**). In contrast to the batch culture, qPCR on the feces revealed that only one of the three *E. coli* phages (phage P10) was detectable at high levels in the feces (**S3B Fig**). Levels of phage P10 remained stable over time at approximately 10^8^ copies/g feces, whereas phage P3 was only detectable on day 1 and 2 p.c. and then decreased below the detection limit of the qPCR assay. Phage P17 was also only detectable at early time points in half of the mice (3 out of 6) at very low abundances (10^5^ copies/g feces – 10^6^ copies/g feces), close to the detection limit of the qPCR. On the other hand, all three *E. faecalis* phages were detectable in feces via qPCR until day 7 of the experiment (**S3C Fig**). Str2 and Str6 were highly abundant (10^7^ copies/g feces – 10^9^ copies/g feces), whilst abundances of Str1 was lower but still above the detection limit in most of the mice.

To investigate if phage treatment causes inflammatory changes in the gut of treated mice, we quantified fecal levels of lipocalin-2 (LCN2), an inflammation marker (zit) by ELISA. No changes in LCN2 levels were observed over time or between the treatment groups (data not shown).

Additionally, we also quantified short-chain fatty acid (SCFA) levels in feces over the course of the experiment in order to assess potential collateral effects of the phage treatments on bile acid metabolism (**S4 Fig**). For all measured SCFAs (2-Methylbutyric acid, Acetic acid, Butyric acid, Isobutyric acid, Isovaleric acid, Lactic acid, Propionic acid, Valeric acid), no difference was observed in treatment versus control groups.

### Phage cocktails targeting *E. coli* and *E. faecalis* leads to decreased colonization resistance of OMM^14^ mice against *S.* Tm

To test the effect of phages on colonization resistance, we used an avirulent *Salmonella enterica* serovar Typhimurium strain (*S*. Tm^avir^), which colonizes the gut but does not induce inflammation due to the lack of functional type III secretion systems 1 and 2 (19). OMM^14^ mice (n = 6-8) were orally infected with *S*. Tm^avir^ (1×10^7^ CFU) directly followed by either the 3Φ^Mt1B1^ or 3Φ^KB1^ phage cocktail (1×10^7^ PFU of each phage in PBS) or PBS control without phage (**Fig 3A**). Fecal samples were collected one and two days post infection (p. i.) with *S*. 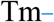. As confirmed in the previous experiment, *E. coli* Mt1B1 and *E. faecalis* KB1 loads significantly decreased at day one p.c. by about one (*E. coli*) and two (*E. faecalis*) orders of magnitude (**Fig 3B** and **3C**). Phages were detectable in the gut for two days, with loads ranging between 4.2×10^4^ PFU/g feces and 7.07×10^7^ PFU/g feces. (**Fig 3D)**. *S*. Tm loads on day one p.i. showed no difference between the control group and any of the phage treatment groups (**Fig 3E** and **3F**). Strikingly, at day two p. i., *S*. Tm loads were significantly increased in mice treated with phage cocktails compared to the control groups (**Fig 3E**). This was more pronounced in the case of mice treated with 3Φ^KB1^, in which the *E. faecalis*-specific cocktail was associated with an increase in *S*. Tm loads by two orders of magnitude (**Fig 3F**).

**Fig. 3:**
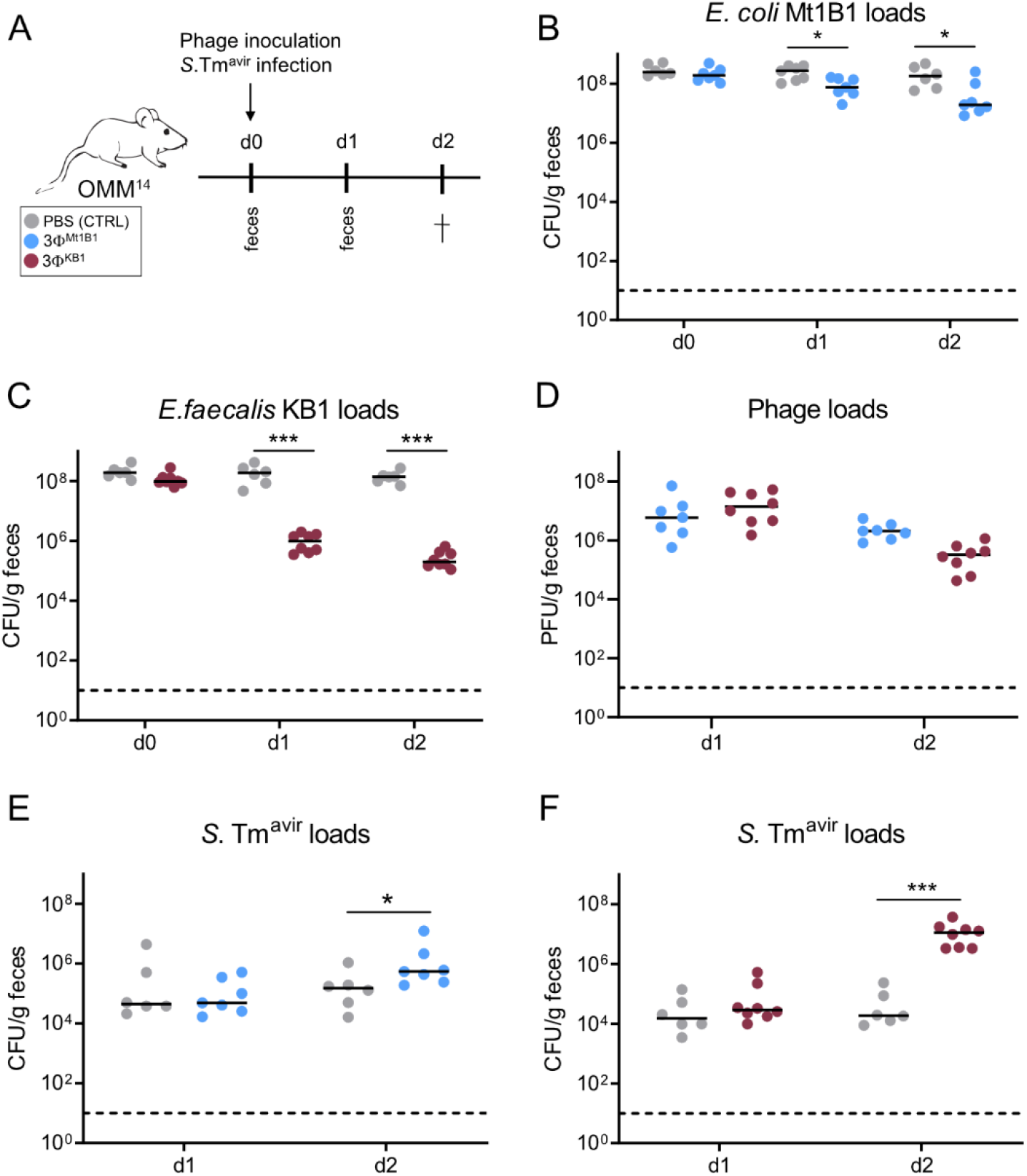
Treatment with phage cocktails targeting *E. coli* and *E. faecalis* leads to increased *S*. Tm loads after infection of OMM^14^ mice. **(A)** Experimental setup, mice stably colonized with the OMM^14^ community were infected with *S*. Tm^avir^ (5×10^7^ CFU) and directly after challenged orally with phage cocktails 3Φ^Mt1B1^ or 3Φ^KB1^ (10^7^ PFU per phage) or PBS as control. Feces were taken at day one and two p.c. and mice were sacrificed at day two p.c.. **(B)** *E. coli* Mt1B1 and **(C)** *E. faecalis* KB1 loads (CFU/g feces) were determined in feces by plating. **(D)** Phage loads (PFU/g feces) were determined by spot assays. **(E, F)** *S*. Tm^avir^ loads at day 1 and 2 after phage challenge (= day 1 and 2 post infection (p.i.)). Statistical analysis was performed using the Mann-Whitney Test (* p<0.05, ** p<0.01, *** p<0.001, N=6-8). Each dot represents one mouse, black lines indicate median, dotted lines indicate limit of detection.

We next tested if phages which do not amplify in OMM^14^ mice due to the lack of host strains would alter the susceptibility to *S*. Tm infection. We gavaged OMM^14^ mice with either a single phage strain (vB_SauP_EBHT, 1×10^7^ PFU in 100μL PBS), targeting *Staphylococcus aureus* (strain EMRSA-15) or 100μL PBS and infected them with *S*. Tm^avir^ **(S5A Fig**). We had verified previously that vB_SauP_EBHT did not target any of the OMM^14^ strains *in vitro*. No significant difference in *S*. Tm^avir^ loads were observed between the control group (p = 0.42) and those treated with phage vB_SauP_EBHT **(S5B Fig**), suggesting that phage treatment *per se* does not alter colonization resistance. Phage loads were determined via spot assays on *S. aureus*, but were only detectable in low numbers at day 1 p.c. **(S5E Fig)**.

### Treatment with phages targeting *E. faecalis* KB1 facilitates development of *S.* Tm-induced colitis in OMM^14^ mice

The observed results proposed to test whether phage-mediated disruption of colonization resistance in OMM^14^ mice would also enhance symptoms of *S*. Tm-induced colitis. Therefore, OMM^14^ mice were infected with *S*. Tm^wt^ (1×10^7^ CFU) and additionally with either 3Φ^Mt1B1^ or 3Φ^KB1^ phage cocktail (1×10^7^ PFU of each phage in PBS or PBS control; **Fig 4A**). Fecal samples were collected one and two days post infection (p. i.). Phage levels were comparable to the previous *in vivo* experiments and stayed stable until day four p. c. despite the inflammation in the gut (**S6A Fig**). In this experiment, loads of *E. coli* Mt1B1 significantly decreased at day one p. c. and *E. faecalis* KB1 loads were significantly decreased at day 1- 4 p. c. (**Fig 4B** and **4C**). *S*. Tm^wt^ loads showed a significant increase on day two and three p. c. (**Fig 4D**) in the group treated with *E. faecalis* 3Φ^KB1^ phage cocktail compared to the control group. Strikingly, in this group, *S*. Tm inflammation was enhanced as determined by increased fecal lipocalin-2 levels at day 3 p. i.. No difference was found at day 4 p. i. both in lipocalin-2 levels and cecal histology (**Fig 4E** and **4F**). In contrast, no change in *S*. Tm inflammation was observed in the group treated with *E. coli* 3Φ^Mt1B1^ phage cocktail. The absolute abundances of all members of the OMM^14^ consortium as well as the phages was determined by qPCR (**S6C Fig**). In general, overall bacterial levels were stable until day three, followed by a strong variation in abundance in all treatment groups on day four (**S6C Fig**). This effect is likely due to coinciding severe *S*. Tm-induced gut inflammation. In conclusion, we verified that phage treatment accelerates not only pathogen invasion but also disease onset in case of the 3Φ^KB1^ phage cocktail.

**Fig. 4:**
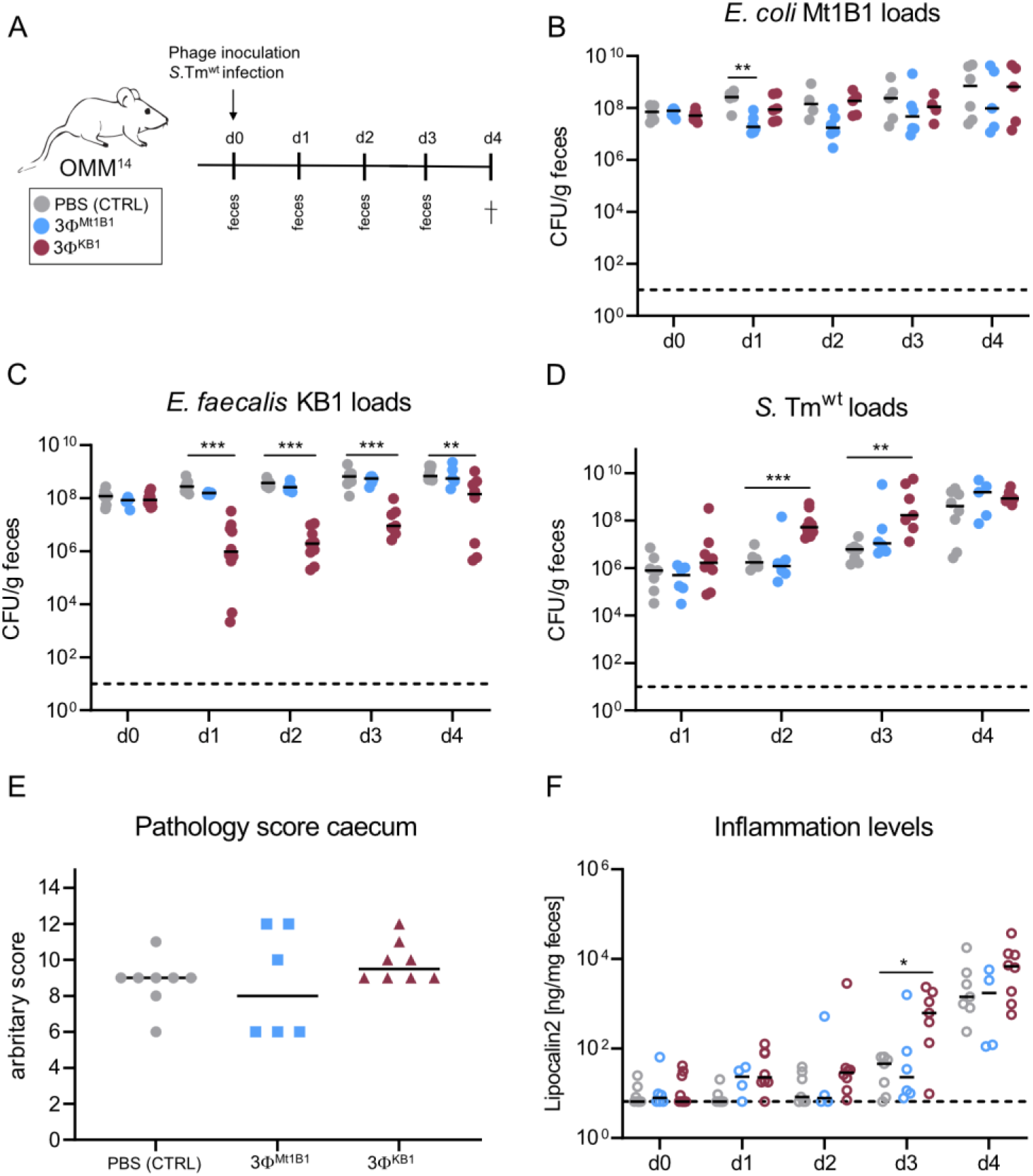
Treatment with phage cocktail targeting *E. faecalis* facilitates *S*. Tm induced colitis in OMM^14^ mice. **(A)** Experimental setup, mice stably colonized with the OMM^14^ community were challenged orally with phage cocktails 3Φ^Mt1B1^ or 3Φ^KB1^ or PBS as control (10^7^ PFU per phage) and infected with *S*. Tm^wt^ (5×10^7^ CFU). Feces were taken at day 1-3 p.c. and mice were sacrificed at day 4 p.c. **(C)** *E. coli* Mt1B1, **(B)** *E. faecalis* KB1 and **(D)** *S*. Tm^wt^ loads (CFU/g) were determined in feces by plating. **(E)** Histopathological score of the cecum at day 4. **(F)** Inflammation levels were determined by measuring the Lipocalin-2 (ng/mg feces) levels in the feces, utilizing a specific ELISA. Statistical analysis was performed using the Mann-Whitney Test (* p<0.05, ** p<0.01, *** p<0.001, N=6-8). Each dot represents one mouse, black lines indicate median, dotted lines indicate limit of detection.

### Phage cocktails impair colonization resistance independently of changing abundance of protective bacteria

To investigate whether phages could also have a long-term impact on colonization resistance, we inoculated groups of OMM^14^ mice with 3Φ^Mt1B1^ or 3Φ^KB1^ phage cocktails (or PBS control) and then infected them with *S*. Tm^avir^ (1×10^7^ CFU) on day seven p.c.. Fecal samples were collected one and two days post infection (p. i.) (**Fig 5A**), where no significant differences in *E. coli* Mt1B1 or *E. faecalis* KB1 loads were observed anymore between the groups (**Fig 5B** and **5C**) but phages were still present at steady numbers in the feces (**Fig 5D**). Further, overall OMM^14^ composition had reverted to the state before treatment (**S3A Fig**). Interestingly, *S*. Tm^avir^ loads at day 8 and 9 p.c. (corresponding to day one and two p.i.) were significantly increased in both phage-treated groups (**Fig 5E**). This suggests that phages can also impair colonization resistance without detectable impact on their target bacteria or overall composition of the microbiota.

**Fig. 5:**
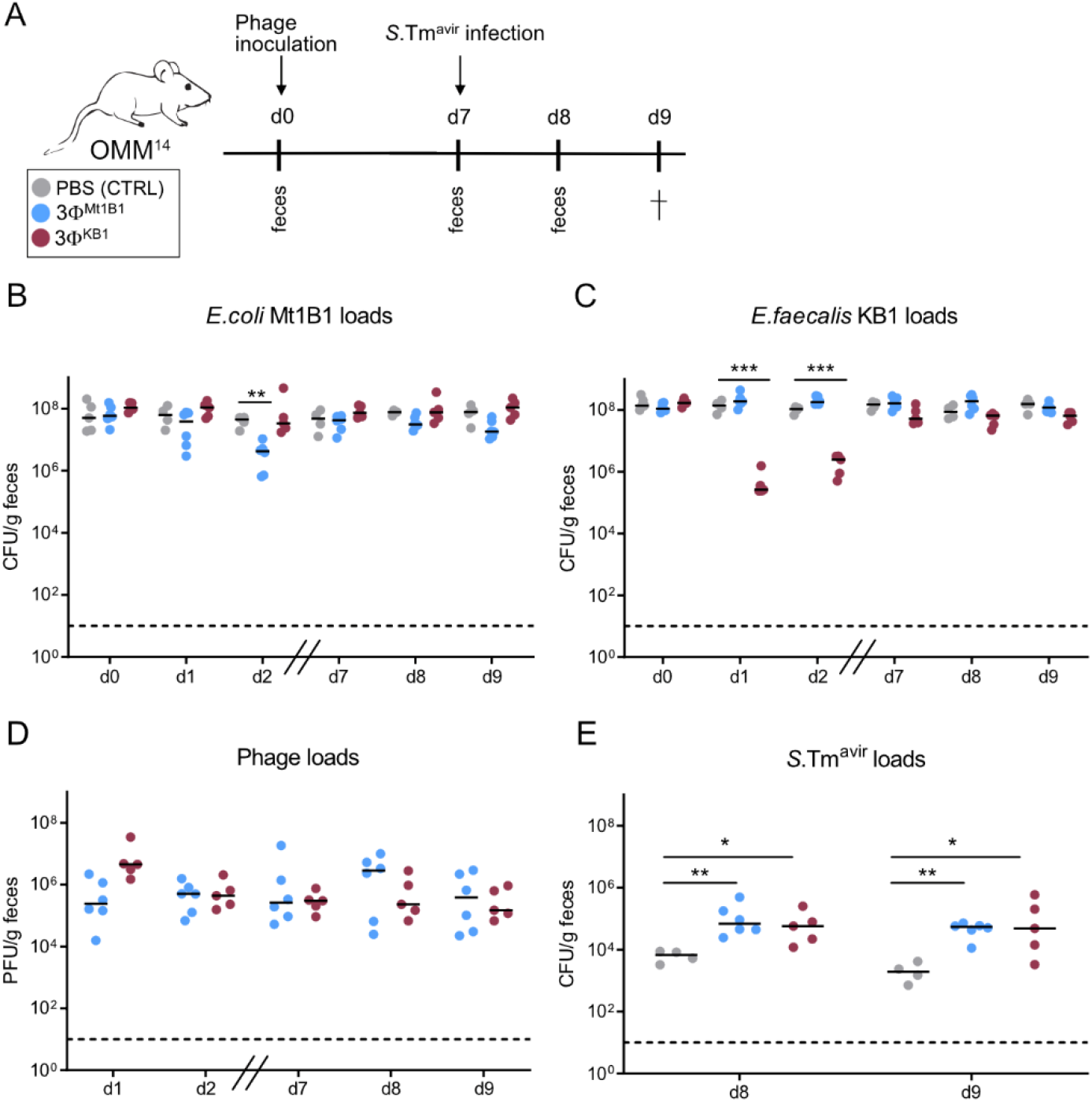
Phages impair colonization resistance at late time points independently of changing abundance of protective bacteria. **(A)** Experimental setup, mice stably colonized with the OMM^14^ community were challenged orally with phage cocktails 3Φ^Mt1B1^ or 3Φ^KB1^ (10^7^ PFU per phage) or PBS as control and infected with *S*. Tm^avir^ (5×10^7^ CFU) at day 7 p.c.. Feces were taken at day 1, 2, 7, 8 and 9 p.c. and mice were sacrificed at day 9 p.c. (corresponding to day two p.i.). **(B)** *E. coli* Mt1B1 and **(C)** *E. faecalis* KB1 loads (CFU/g) were determined in feces by plating. **(D)** Phage loads (PFU/g feces) of 3Φ^Mt1B1^ phages and 3Φ^KB1^phages were determined by spot assays in fecal samples at different time points p.c.. **(E)** *S*. Tm^avir^ loads at day 8 and 9 p.c. (= day 1 and 2 p.i.). Statistical analysis was performed using the Mann-Whitney Test (* p<0.05, ** p<0.01, *** p<0.001, N=4-6). Each dot represents one mouse, black lines indicate median, dotted lines indicate limit of detection.

## Discussion

Phages are the most prevalent viruses in the gut (27, 28) and coexist with their host bacteria over long periods (17). They play an important role in shaping composition, dynamics (29) and evolution (30) of microbial communities (31). There is a growing recognition for the therapeutic use of phages against intestinal colonization by multi-drug resistant strains and bacteria causally involved in the pathogenesis of inflammatory bowel diseases (32). Nevertheless, the functional impact of phages on bacterial communities is not fully understood. Using mice stably associated with a community of 14 commensal bacterial species, we show that virulent phages targeting specific strains in the community can impair colonization resistance and increase susceptibility of mice to oral infection by a human intestinal pathogen.

Phage infection of *E. coli* and *E. faecalis* in OMM^14^ mice can be roughly divided into two phases: A transient reduction of the bacterial target population in the acute infection phase (1-4 days p.c.) paralleled by high initial phage loads. Thereafter, as the second phase, phages and host bacteria co-existed until day 7 p. c.. Previous work demonstrated that co-existence is in part explained by phage-inaccessible sites in the mucosa, which serve as a spatial refuge for part of the host bacterial population (17). In addition, bacterial hosts differentially express genes in the gut that foster phage-bacteria coexistence (33). Phage-specific qPCR analysis allowed us to quantify individual phages. This revealed that the abundance of the three different phages in each cocktail differed markedly in the feces. For example, *E. coli* phage P10 dominated at all time points while loads of phage P3 and P17 decreased rapidly after inoculation below the limit of detection. Interestingly, abundance of *E. faecalis* phages was also different, but all three phages were detectable until seven days p. c.. Different abundances of phages may be due to burst size, stability of the co-existence and perhaps selection of phage-resistant mutants, leading to different abundances and abilities of the individual phages to infect and amplify.

Phage treatment impaired CR against *S*. Tm in the acute phage infection phase. CR in the OMM^12^ model is mediated by competition for substrates including C5/C6 sugars, oxygen and electron acceptors for anaerobic respiration (21), and OMM^12^ mice lacking *S*. Tm competitors *E. coli* Mt1B1 or *E. faecalis* KB1 exhibit loss of CR. Thus, we reason that transient reduction of these two species caused by phage treatment leads to a short-term increase in substrate availability and weakening of the targeted strains opening a window for pathogen invasion. In support of this idea, the degree of reduction of the target population in the acute phase equals the increase in pathogen loads. Besides phage-induced changes in the microbiota, antibiotic-mediated disruption of the microbiota transiently increases levels of free sugars which can be exploited by pathogens (34). Moreover it was shown, that a high-fat diet increasing bile acid release and microbiota changes can also facilitate pathogen invasion (5).

Surprisingly, mice also displayed impaired CR at day 7 post phage challenge, when phages and their target strains stably co-existed in the gut. This hints to mechanisms also acting independently on the reduction of the loads of competing bacteria. Phages may decrease the target population locally, enabling *S*. Tm to invade. In addition, the presence of phages might induce transcriptional changes of the host strains leading to increased resistance against phage infection. These may include expression of anti-phage systems and reprogramming of bacterial metabolism (35) and phage receptors (33). Such potential changes may directly or indirectly affect CR against *S*. Tm. Further work is needed to clarify this interrelation.

Using a wildtype *S*. Tm strain which triggers gut inflammation within 3 days of infection in gnotobiotic mice (21), we observed impaired CR and a faster onset of gut inflammation when mice were treated with the *E. faecalis* cocktail 3Φ^KB1^. However, the 3Φ^Mt1B1^ cocktail had no effect on the course of *S*. Tm^Wt^ infection. This differential effect of phages on avirulent and the wildtype *S*. Tm infection might be due to utilization of different niches of both, *S*. Tm and *E. coli* in the normal versus inflamed intestine (36).

An important selling point of phage therapy is that phage cocktails act in species-specific manner and do not disrupt the overall (bacterial) microbiome composition (10, 37, 38). In our work, we confirm that the phage cocktails specifically targeting *E. coli* Mt1B1 and *E. faecalis* KB1 do not affect the overall bacterial community structure. Even though, we demonstrate that phage cocktails can create a window of opportunity for pathogen invasion, in this case human pathogenic *S*. Tm, if commensals relevant for mediating CR are specifically targeted. Based on our data, we reason that one possible future application of phage cocktails is to facilitate strain-replacement strategies – and mediate exchange of harmful bacteria by beneficial strains in the human microbiome (39, 40).

In conclusion, we show that phage cocktails targeting strains that are functionally important for mediating protection against pathogens can open up niches for pathogens (and potentially also non-pathogenic bacteria) to invade the gut. We reason that phage ingestion in conjunction with pathogen contaminated food or water sources could facilitate pathogen invasion and may therefore be risk factors of human *Salmonella* infection. Coliphage content in water ranges from 8×10^4^ PFU/100mL in wastewater to 30 PFU/100mL for river source waters (41), which might be enough to target protective bacteria in the gut. Eventually, future work will be needed to clarify if therapeutic application of phage cocktails against multidrug resistant *E. coli* or *E. faecalis* strains may also impair CR in the human host and should therefore be handled with caution.

## Methods

### Strains and culture conditions

The following strains were used in this study: *Enterococcus faecalis* KB1 (DSM 32036), *Bifidobacterium animalis* YL2 (DSM 26074), *Acutalibacter muris* KB18 (DSM 26090), *Muribaculum intestinale* YL27 (DSM 28989), *Flavonifractor plautii* YL31 (DSM 26117), *Enterocloster clostridioformis* YL32 (DSM 26114), *Akkermansia muciniphila* YL44 (DSM 26127), *Turicimonas muris* YL45 (DSM 26109), *Clostridium innocuum* I46 (DSM 26113), *Bacteroides caecimuris* I48 (DSM 26085), *Limosilactobacillus reuteri* I49 (DSM 32035), *Blautia coccoides* YL58 (DSM 26115), *Escherichia coli* Mt1B1 (DSM 28618) (26), *Extibacter muris* JM40 (DSM 28560) (25), *S*. Tm^wt^ SL1344 (SB300) (42), *S*. Tm^avir^ M2707 (43), *Staphylococcus aureus* RG2 (DSM 104437).

OMM^14^ cultures were prepared from individual glycerol cryostocks in a 10 ml culture and subculture in cell culture flasks (flask T25, Sarstedt) previous to all *in vitro* experiments. Cultures were incubated at 37°C without shaking under strictly anaerobic conditions (gas atmosphere 7% H_2_, 10% CO_2_, 83% N_2_). OMM^14^ bacterial cultures were grown in anaerobic medium (AF medium, : 18 g.l^−1^ brain-heart infusion (Oxoid), 15 g.l^−1^ trypticase soy broth (Oxoid), 5 g.l^−1^ yeast extract, 2.5 g.l^−1^ K_2_HPO_4_, 1 mg.l^−1^ haemin, 0.5 g.l^−1^ D-glucose, 0.5 mg.l^− 1^ menadione, 3% heat-inactivated fetal calf serum, 0.25 g.l^−1^ cysteine-HCl × H_2_O).

For mouse infection experiments, *S*. Tm strains were grown on MacConkey agar plates (Oxoid) containing streptomycin (50 μg/ml) at 37 °C. One colony was re-suspended in 3 ml LB containing 0.3 M NaCl and grown 12 h at 37 °C on a wheel rotor. A subculture (1:20 dilution) was prepared in LB with 0.3 M NaCl and incubated for 4 h. Bacteria were washed in ice-cold sterile PBS, pelleted and re-suspended in PBS for gavage.

### Spot assays

1 ml of an exponentially growing bacterial culture, containing the phage host, was applied on an EBU agar plate, containing Evans Blue (1%) and Fluorescein sodium salt (1%) to visualize bacterial lysis, which is indicated by a color change to dark green. Excess liquid was removed and the plate was dried under the laminar airflow cabinet for 15 minutes. Phage lysates were serially diluted in sterile PBS and 5 μl of each dilution was spotted on the bacterial lawn. The plate was incubated over night at 37°C. The next day, clear plaques and lysis of the bacteria could be detected. Plaques were counted to quantify plaque forming units (PFU).

### Isolation of *E. faecalis* KB1 phages from sewage water

For isolation of phages specific for *Enterococcus faecalis* KB1, sewage water from different sources was filtered (0.22 μm) and mixed with an equal volume of 2 × LB media. Next, bacterial overnight culture was added with a final dilution of 1/100 and incubated overnight at 37°C without shaking. The next day, the cultures were centrifuged (6,000 × g, 10 min), filtered (0.22 μm) and spotted in serial dilution on a lawn of *E. faecalis* KB1 on EBU plates followed by another incubation at 37 °C overnight. If clear plaques were visible, individual plaques were picked and diluted in 100 μl SM buffer (100 mM NaCl, 8.1 mM MgSO_4_ × H_2_O, 50 mM Tris-HCL (pH 7.5)), added to a fresh bacterial subculture in 10 ml LB medium and incubated overnight. Centrifugation and filtration of this culture resulted in a sterile phage suspension which was stored at 4°C and used for the *in vitro* and *in vivo* experiments.

### Generation of phage cocktails

100 μl of purified and sterile phage suspension was added to 10 ml of LB media and incubated with 100 μl of overnight culture of their respective bacterial host overnight at 37°C. The next day, the cultures were centrifuged (6,000 × g, 10 min), filtered (0.22 μm) and spotted in serial dilution on a lawn of their respective bacterial host on EBU plates followed by another incubation at 37°C overnight. The next day, plaque forming units were calculated for the phage suspensions 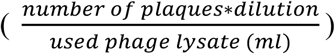 and the phages were mixed together in a concentration of 1×10^7^ PFU/100μl each. The following phages were used in this study: Mt1B1_P3, Mt1B1_P10, Mt1B1_P17 (17), vB_efaS_Str1 (vB_EfaS_Strempel1, DSM 110103), vB_efaP_Str2 (vB_EfaP_Strempel2, DSM 110104), vB_efaS_Str6 (vB_EfaS_Strempel6, DSM 110108), vB_SauP_EBHT (DSM 26856).

### Phage genome sequencing and analysis

The three phages targeting *E. coli* Mt1B1 were sequenced and analyzed as described previously (17). Shortly, sequencing was performed using Illumina MiSeq Nano. Assembly was performed using a workflow implemented in Galaxy-Institut Pasteur and phage termini were determined by PhageTerm. The three *E. faecalis* KB1 phages were also sequenced using Illumina MiSeq Nano and assembled with SPAdes (3.12.0) (44). Annotation was performed using PROKKA (45) and taxonomic classification was performed using VICTOR (46) as previously described (47).

### Growth measurements

Bacterial growth was measured in 96 well plates (Sarstedt TC-cellculture plate) using an Epoch2 plate reader (GenTech). Inocula were prepared from an overnight culture and subculture and diluted in AF medium (20) to 0.01 OD_600_. To determine phage growth and bacterial lysis, 150 μl of bacterial culture was added to each well, followed by either 10 μl of PBS as a control or 10 μl of sterile phage lysate with a defined concentration of active phages (PFU/ml), resulting in a MOI of 0.01. During continuous measurements, the plate was heated inside the reader to 37°C and a 30 second double orbital shaking step was performed prior to every measurement. Measurements took place every 15 minutes for a duration of 20 hours in total.

### OMM^14^ community cultures

OMM^14^ communities were cultured as previously described (20). Shortly, monoculture inocula were prepared from an overnight culture and subculture and were diluted to OD_600_ 0.1 in AF medium. Following, the community inoculum with equivalent ratios of all 14 strains was generated from this dilution. The inoculum was distributed to 24 well plates, thereby diluting the inoculum 1:10 with 0.9 ml AF medium, resulting in a starting OD_600_ of 0.01 (time = 0h). 24-well plates were incubated at 37°C without shaking under anaerobic conditions. Every 24 h, samples were taken for qPCR analysis and spot assays, and cultures were diluted 1:100 in AF medium. Phages were added to the community 12h after the third dilution, in a concentration of 1*10^6^ PFU per phage in 10 μl PBS.

### Animal experiments

Germ-free mice were inoculated with OMM^12^ cryostock mixtures as described (26) and individual frozen stocks of *E. muris* and *E. coli* Mt1B1. Stocks were thawn in 1% Virkon S (V.P. Produkte) disinfectant solution and used for inoculation of germ-free C57Bl/6 mice in a flexible film isolator. Mice were inoculated twice (72 h apart) with the bacterial mixtures and the two single stocks by gavage (50 μl orally, 100 μl rectally). Mice were housed and bred under germfree conditions in flexible film isolators (North Kent Plastic Cages). For experiments, mice were transferred into isocages (IsoCage P system, Tecniplast). Mice were supplied with autoclaved ddH_2_O and autoclaved Mouse-Breeding complete feed for mice (Ssniff) *ad libitum*. For the following experiments, only mice starting from generation F1 were used, to ensure stable colonization of the consortium.

For all experiments, female and male mice between 6-20 weeks were used and animals were assigned to experimental groups to match sex and age. Mice were kept in groups of 2-6 mice/cage during the experiment. All animals were scored twice daily for their health status. Phage cocktails containing the same concentration of phages as for the *in vitro* experiments (10^7^ PFU per phage in 100 μl PBS) were administered by oral gavage. Mice were infected with *S*. Tm by oral gavage with 50 μl of bacterial suspension (approximately 5×10^7^ CFU). All mice were sacrificed by cervical dislocation. Feces and cecal content were weighed and dissolved in 500 μl sterile PBS and *S*. Tm and *E. coli* Mt1B1 loads were determined by plating in several dilutions on MacConkey agar (Oxoid) supplemented with respective antibiotics (vancomycin 7.5 μg/ml for *E. coli*, streptomycin 50 μg/ml for *S*. Tm). *E. faecalis* KB1 loads were determined by plating in several dilutions on BHI agar (Oxoid), supplemented with polymycin B 50 μg/ml. 100 μl of dissolved intestinal content were sterile filtered using Centrifuge Tube Filter (0.22 μm, Costar Spin-X) and spotted in serial dilutions in PBS on a lawn of *E. coli* and *E. faecalis* on EBU agar plates to determine the phage loads. Lipocalin-2 was quantified from supernatant of frozen cecal content. For metabolomics, samples were directly snap frozen in liquid N_2_ and stored at −80°C until further processing. Samples for DNA extraction were stored at −20°C after weighing.

### DNA extraction from intestinal content

gDNA extraction was performed using a phenol-chloroform based protocol as described previously (48). Briefly, fecal pellet or cecal content was resuspended in 500 μl extraction buffer (200 mM Tris-HCl, 200 mM NaCl, 20 mM EDTA in ddH_2_O, pH 8, autoclaved), 210 μl 20 % SDS and 500 μl phenol:chloroform:isoamylalcohol (25:24:1, pH 7.9). Furthermore, 500μl of 0.1 mm-diameter zirconia/silica beads (Roth) were added. Bacterial cells were lysed with a bead beater (TissueLyser LT, Qiagen) for 4 min, 50 Hz. After centrifugation (14,000 × g, 5 min, RT), the aqueous phase was transferred into a new tube, 500 μl phenol:chloroform:isoamylalcohol (25:24:1, pH 7.9) were added and again spun down. The resulting aqueous phase was gently mixed with 1 ml 96 % ethanol and 50 μl of 3 M sodium acetate by inverting. After centrifugation (30 min, 14,000 × g, 4 °C), the supernatant was discarded and the gDNA pellet was washed with 500 μl ice-cold 70 % ethanol and again centrifuged (14,000 × g, 4 °C; 15 min). The resulting gDNA pellet was resuspended in 100 μl Tris-HCL pH 8.0. Subsequently, gDNA was purified using the NucleoSpin gDNA clean-up kit (Macherey-Nagel) and stored at −20°C.

### Quantitative PCR for bacteria and phages

Quantitative PCR was performed as described previously (for the OMM^12^ strains and *E. coli* Mt1B1 see (19), for *E. muris* JM40 see (25)). Standard curves using linearized plasmids containing the 16S rRNA gene sequence of the individual strains were used for absolute quantification of 16S rRNA gene copy numbers of individual strains.

Probe and primer design for the phages was done using the software PrimerExpress. 1500 bp of either the phage tail fiber gene or the phage major capsid protein gene were used as a template gene instead of the bacterial 16S rRNA gene to design probes, primers and plasmids (**Table S1**). Absolute quantification was conducted as for the bacterial qPCR (19).

### Lipocalin-2 quantification

Lipocalin-2 levels in feces and cecal content were determined by an enzyme-linked immunosorbent assay (ELISA) kit and protocol from R&D Systems (DY1857, Minneapolis, US), following manufacturer’s instructions. Absorbance was measured at OD_405_.

### Targeted short chain fatty acid (SCFA) measurement

Feces and gut content (approximately 20 mg) for metabolomic profiling were freshly sampled, weighed in a 2 ml bead beater tube (CKMix 2 mL, Bertin Technologies, Montigny-le-Bretonneux, France) filled with ceramic beads (1.4 mm and 2.8 mm ceramic beads i.d.) and immediately snap frozen in liquid nitrogen. The following procedure and measurement was performed at the BayBioMS (TU Munich). 1 mL methanol was added and the sample was homogenized by bead beating using a bead beater (Precellys Evolution, Bertin Technologies) supplied with a Cryolys cooling module (Bertin Technologies, cooled with liquid nitrogen) 3 times each for 20 seconds with 15 seconds breaks in between, at a speed of 10.000 rpm. After centrifugation of the suspension (10 min, 8000 rpm, 10°C) using an Eppendorf Centrifuge 5415R (Eppendorf, Hamburg, Germany), the 3-NPH method was used for the quantitation of SCFAs (49). Briefly, 40 μL of the fecal extract and 15 μL of isotopically labeled standards (ca 50 μM) were mixed with 20 μL 120 mM EDC HCl-6% pyridine-solution and 20 μL of 200 mM 3-NPH HCL solution. After 30 min at 40°C and shaking at 1000 rpm using an Eppendorf Thermomix (Eppendorf, Hamburg, Germany), 900 μL acetonitrile/water (50/50, v/v) was added. After centrifugation at 13000 U/min for 2 min the clear supernatant was used for analysis. The measurement was performed using a QTRAP 5500 triple quadrupole mass spectrometer (Sciex, Darmstadt, Germany) coupled to an ExionLC AD (Sciex, Darmstadt, Germany) ultra high performance liquid chromatography system. The electrospray voltage was set to −4500 V, curtain gas to 35 psi, ion source gas 1 to 55, ion source gas 2 to 65 and the temperature to 500°C. The MRM-parameters were optimized using commercially available standards for the SCFAs. The chromatographic separation was performed on a 100 × 2.1 mm, 100 Å, 1.7 μm, Kinetex C18 column (Phenomenex, Aschaffenburg, Germany) column with 0.1% formic acid (eluent A) and 0.1% formic acid in acetonitrile (eluent B) as elution solvents. An injection volume of 1 μL and a flow rate of 0.4 mL/min was used. The gradient elution started at 23% B which was held for 3 min, afterward the concentration was increased to 30% B at 4 min, with another increase to 40% B at 6.5 min, at 7 min 100% B was used which was hold for 1 min, at 8.5 min the column was equilibrated at starting conditions. The column oven was set to 40°C and the autosampler to 15°C. Data aquisition and instrumental control were performed with Analyst 1.7 software (Sciex, Darmstadt, Germany).

### Hematoxylin and eosin staining (HE staining) and histopathological scoring

HE staining was performed as described previously (Herp et al., 2019). Cecal tissue, directly frozen in O.C.T, was cut in 5 μm sections using a cryotome (Leica) and mounted onto Superfrost Plus glass slides (Hartenstein). Sections were dried o/n. and fixed in Wollman solution (95 % ethanol, 5 % acetic acid) for 30 s, washed in flowing tap water (1 min) and rinsed in dH_2_O. Afterwards, slides were incubated in Vectors’s Hämalaun (Roth) for 20 min, washed in flowing tap water (5 min), dipped in de-staining solution (70 % ethanol with 1 % HCl) once, washed again in flowing tap water (5 min) and rinsed in dH_2_O with subsequent rinses in 70 % and 90 % ethanol. Slides were then dipped for 15 s in alcoholic eosin (90 % ethanol) with Phloxin (Sigma-Aldrich), rinsed in dH_2_O followed by dehydration in 90 % ethanol, 100 % ethanol and xylene. Sections were directly mounted with Rotimount (Roth) and dried thoroughly.

Histopathological scoring of cecal tissue was performed as described previously (50). Submucosal edema (0-3), infiltration of polymorphonuclear neutrophils (PMNs) (0-4), loss of goblet cells (0-3) and epithelial damage (0-3) was evaluated and all individual scores were summed up to give a final pathology score: 0-3 no inflammation; 4-8 mild inflammation; 9-13 profound inflammation.

### Generation of a 16S rRNA gene-based phylogenetic tree

The genomes of the twelve strains of the OMM^12^ consortium (51) were accessed via DDBJ/ENA/GenBank using the following accession numbers: CP022712.1, NHMR02000001-NHMR02000002, CP021422.1, CP021421.1, NHMQ01000001-NHMQ01000005, NHTR01000001-NHTR01000016, CP021420.1, NHMP01000001-NHMP01000020, CP022722.1, NHMU01000001-NHMU01000019, NHMT01000001-NHMT01000003, CP022713.1, CP028714, KR364761.1 and annotated using Prokka (default settings) (52). The 16S rRNA sequences of all strains were obtained. These rRNA FASTA sequences were uploaded to the SINA Aligner v1.2.11 (53) to align these sequences with minimum 95% identity against the SILVA database. By this, a phylogenetic tree based on RAxML, GTR Model and Gamma rate model for likelihood was reconstructed. Sequences with less than 90% identity were rejected. The obtained tree was rooted using *midpoint.root()* in the phytools package (54) in R and visualized using iTOL online (55).

### Generation of 16s rRNA gene based tree different *Enterococcus* strains

16S rRNA gene sequences of all tested *Enterococcus* strains and selected reference type strains of the family *Enterococcaceae* were downloaded from NCBI and aligned with Mega-X Version 10.1.5. A Maximum Likelihood tree was constructed using the Tamura-Nei model and the Nearest-Neighbor-Interchange method. The tree was exported in Newick tree format and annotation was done using iTOL online (55).

### Statistical analysis

Statistical details for each experiment are indicated in the figure legends. Mann-Whitney U test and Kruskal-Wallis test were performed using the software GraphPad Prism version 5.01 for Windows (GraphPad Software, La Jolla California USA, www.graphpad.com). P values of less than 0.05 were considered as statistically significant and only those are indicated in the figures (*P<0.05, **P<0.01, ***P<0.001).

## Acknowledgements

This research was supported by Collaborative Research Center 1371 funded by the German Research Foundation (project number 395357507, P14, Z01), the European Research Council (EVOGUTHEALTH; grant no. 865615), DFG-ANR project PhaStGut (STE 1971/11-1), the DFG Priority Programme SPP2330, the German Center for Infection Research (DZIF) and the Center for Gastrointestinal Microbiome Research (CEGIMIR). We are grateful to Saib Hussain and the Germ-free Rodent Facility of the Max von Pettenkofer Institute, LMU Munich for excellent technical support.

## Author contributions

Conceived and designed the experiments: B.S., A.v.S; Performed the experiments: A.v.S., A.S.W., M.S.S., E.W., K.K.; Analyzed the data: A.v.S., K.K., M.G.; B.S. Contributed materials/analysis tools: L.D., E.W., T.C.; Secured funding: B.S.; A.v.S. coordinated the project and wrote the original draft; all authors reviewed and edited the draft manuscript. Correspondence and requests for materials should be addressed to B.S..

