## Supplementary information for "Bacteriophages targeting protective commensals impair resistance against *Salmonella* Typhimurium infection in gnotobiotic mice"

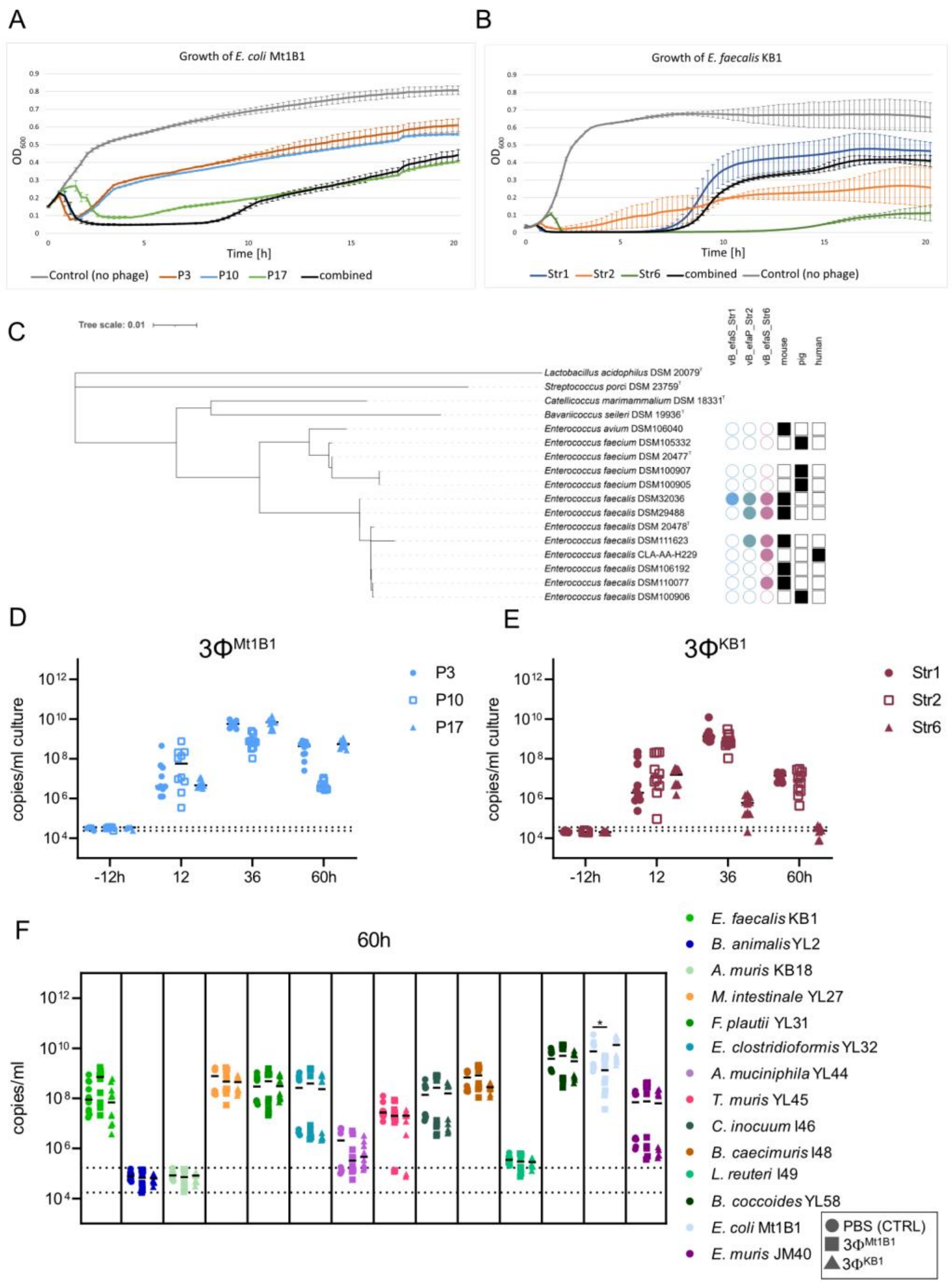


**Fig S1: Characterization of phage cocktails in vitro**

**(A)** Growth curve of *E. coli* Mt1B1 in LB medium (grey) measured under anaerobic conditions at 37°C, challenged with phage P3 (red), P10 (blue), P17 (green) and all three phages combined (black). **(B)** Growth curve of *E. faecalis* KB1 in BHI medium (grey) measured under anaerobic conditions at 37°C, challenged with phage Str1 (blue), Str2 (orange), Str6 (green) and all three phages combined (black). **(C)** Host range of phages Str1, Str2 and Str6 on different *E. faecalis* and *E. faecium* isolates, displayed as a family tree of the bacteria. Filled circles stand for susceptibility, empty circles stand for resistance. **(D)** abundance of single phages of the phage cocktails $3\Phi^{Mt1B1}$ and **(E)** $3\Phi^{KB1}$ in copies per ml batch culture from the experiment shown in Fig.1. Different shapes show the different phages, respectively. **(F)** Community composition 60h after phage addition from experiment shown in Fig.1, absolute abundance of each strain was determined using a strain-specific qPCR and plotted as 16S rRNA copy numbers per ml culture. Statistical analysis was performed using the Mann-Whitney Test comparing the treatment groups (N=10) against the control group (N=10) (* $p<0.05$, ** $p<0.01$, *** $p<0.001$). Each dot represents one well, black lines indicate median, dotted lines indicate detection limit (DTL).

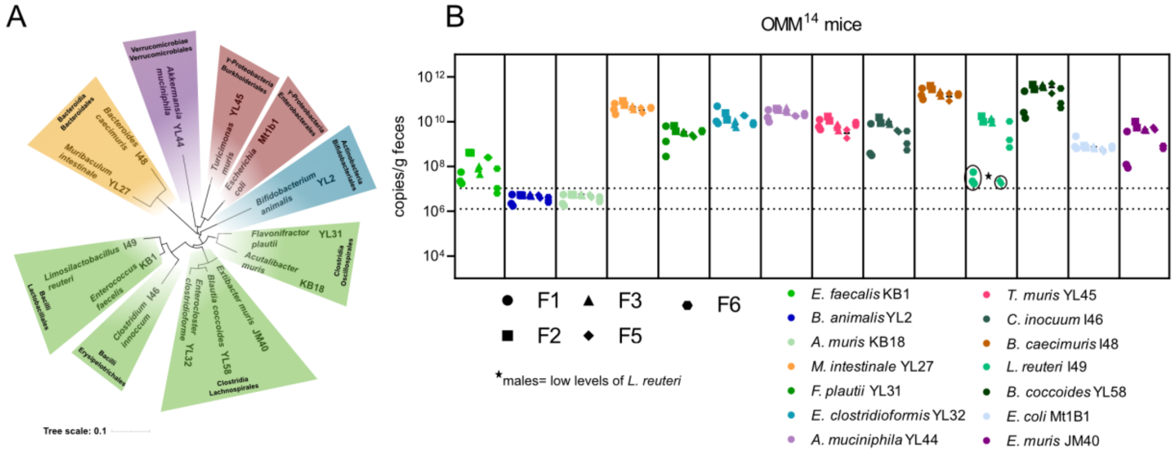

**Fig S2: OMM14 bacterial community**

**(A)** Phylogenetic tree of the OMM14 bacterial community based on 16S rRNA sequences. Different colors represent different phyla. **(B)** Absolute abundance of all 14 bacteria overall several breeding generations (F1, F2, F3, F5 and F6, indicated by different shapes), determined by strain-specific qPCR in 16S rRNA copies per gram feces. Each dot represents one mouse, black line indicates median, dotted lines indicate DTL.

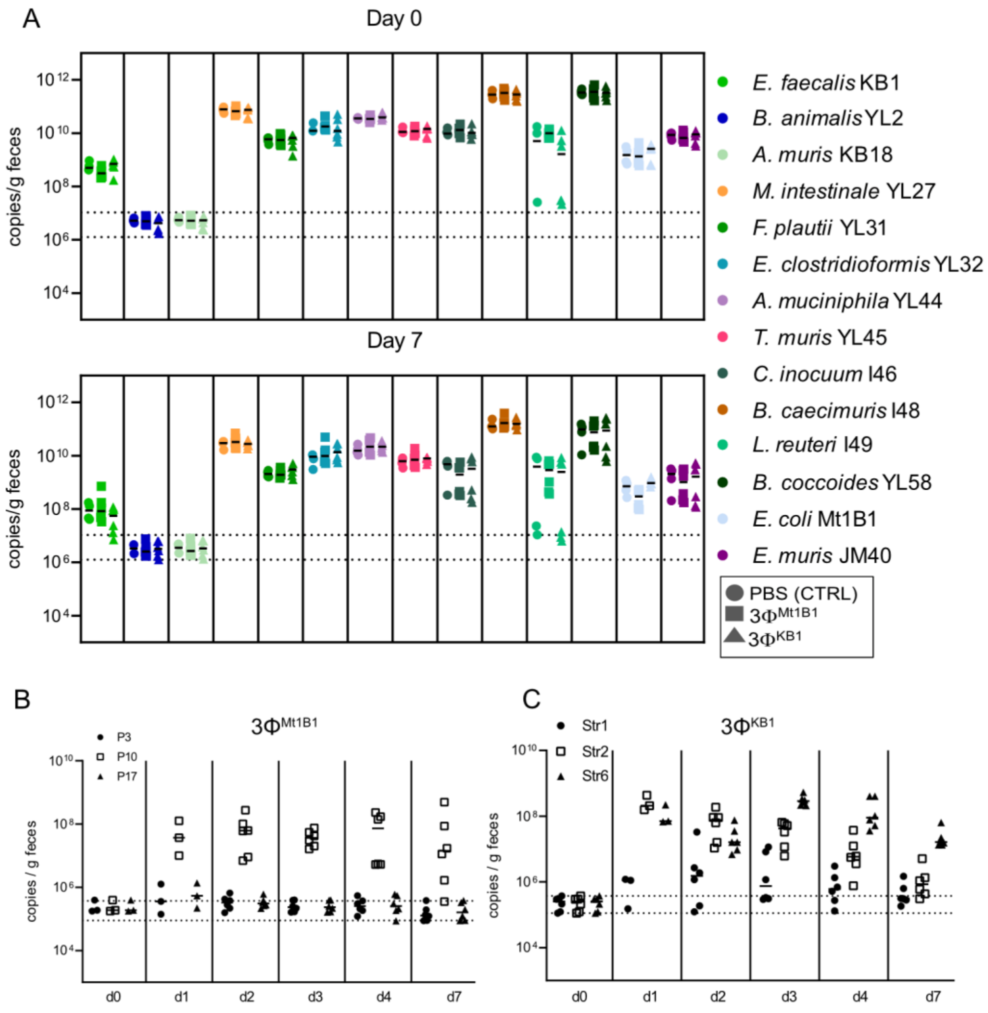


**Fig S3: Abundance of OMM14 community members and single phages in vivo**

**(A)** Absolute abundances of all 14 bacteria from experiment shown in Fig.2, determined by strain-specific qPCR on day 0 and day 7 p. c.. Each color represents one bacterial strain, different shapes represent different experimental groups. **(B)** Abundances of the single phages of the phage cocktails $3\Phi^{Mt1B1}$ and **(C)** $3\Phi^{KB1}$, determined by specific qPCR. Different shapes show different phages. Each dot represents one mouse, black line indicates median, dotted lines indicate DTL.

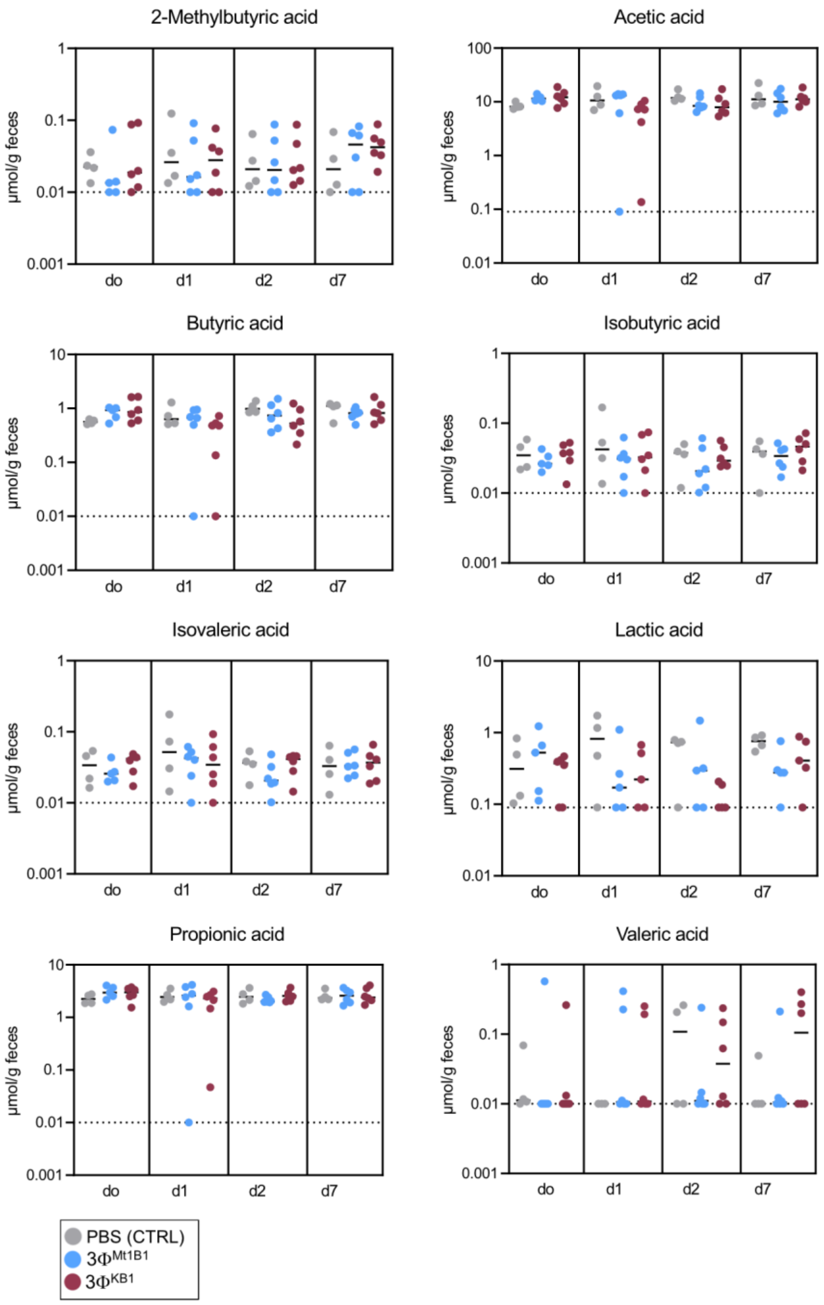


**Fig S4: Short chain fatty acid (SCFA) levels in the mouse gut**

Different SCFAs were measured by quantitative mass spectronomy in fecal samples from the experiment shown in Fig. 2. Different colors represent the different experimental groups, each dot represents one mouse, black line indicates median, dotted lines indicate DTL.

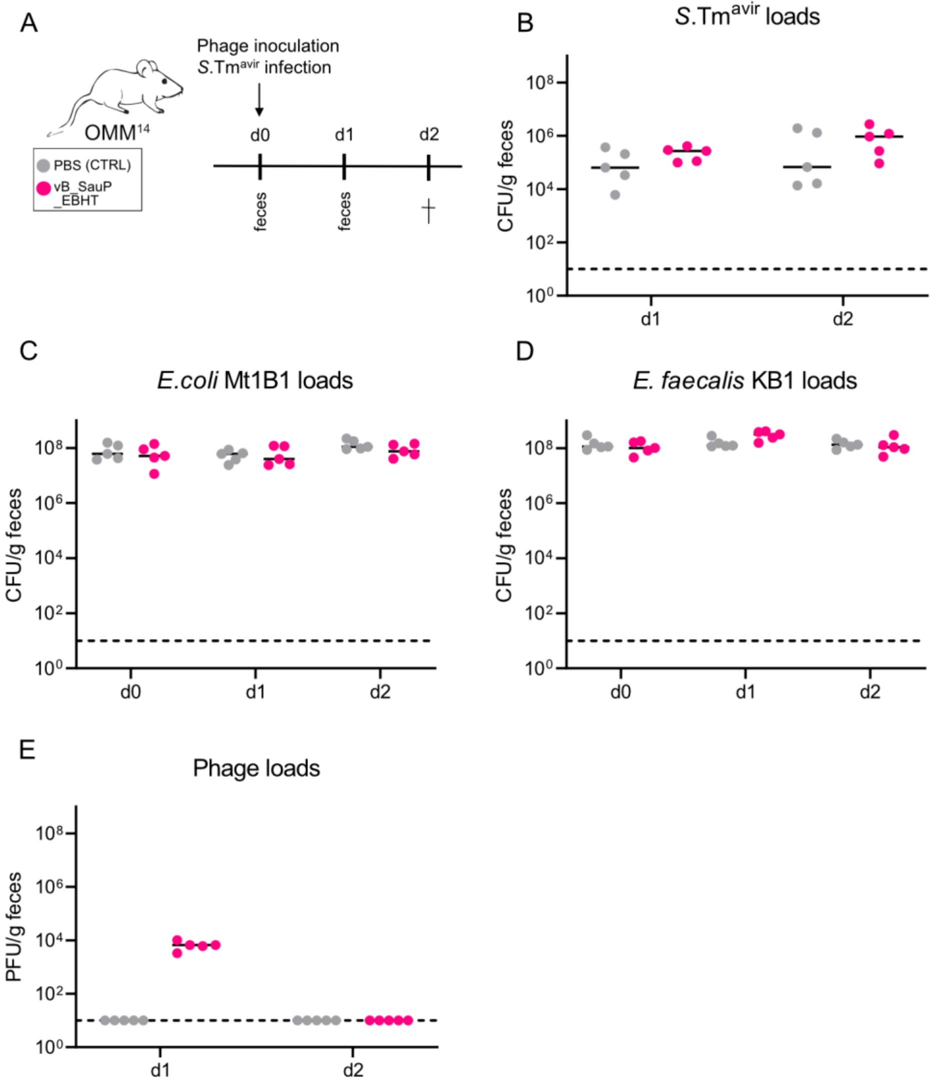


**Fig S5: Treatment with an unspecific phage does not impair the colonization resistance against *S*. Tm$^{avir}$**

**(A)** Experimental setup: mice stably colonized with the OMM$^{14}$ community were challenged orally with phage vB_SauP_EBHT ($10^7$ PFU) or PBS as a control and at the same time challenged with *S.* Tm$^{avir}$ ($5x10^7$ CFU). Feces for plating were taken and mice were sacrificed on day 2 p. i.. **(B)** *S.* Tm*avir* loads, **(C)** *E. coli* Mt1B1 loads and **(D)** *E. faecalis* KB1 loads were monitored via plating. **(E)** phage loads were determined via spot assays on S. aureus. Each dot represents one mouse, black line indicates median, dotted lines indicate DTL.

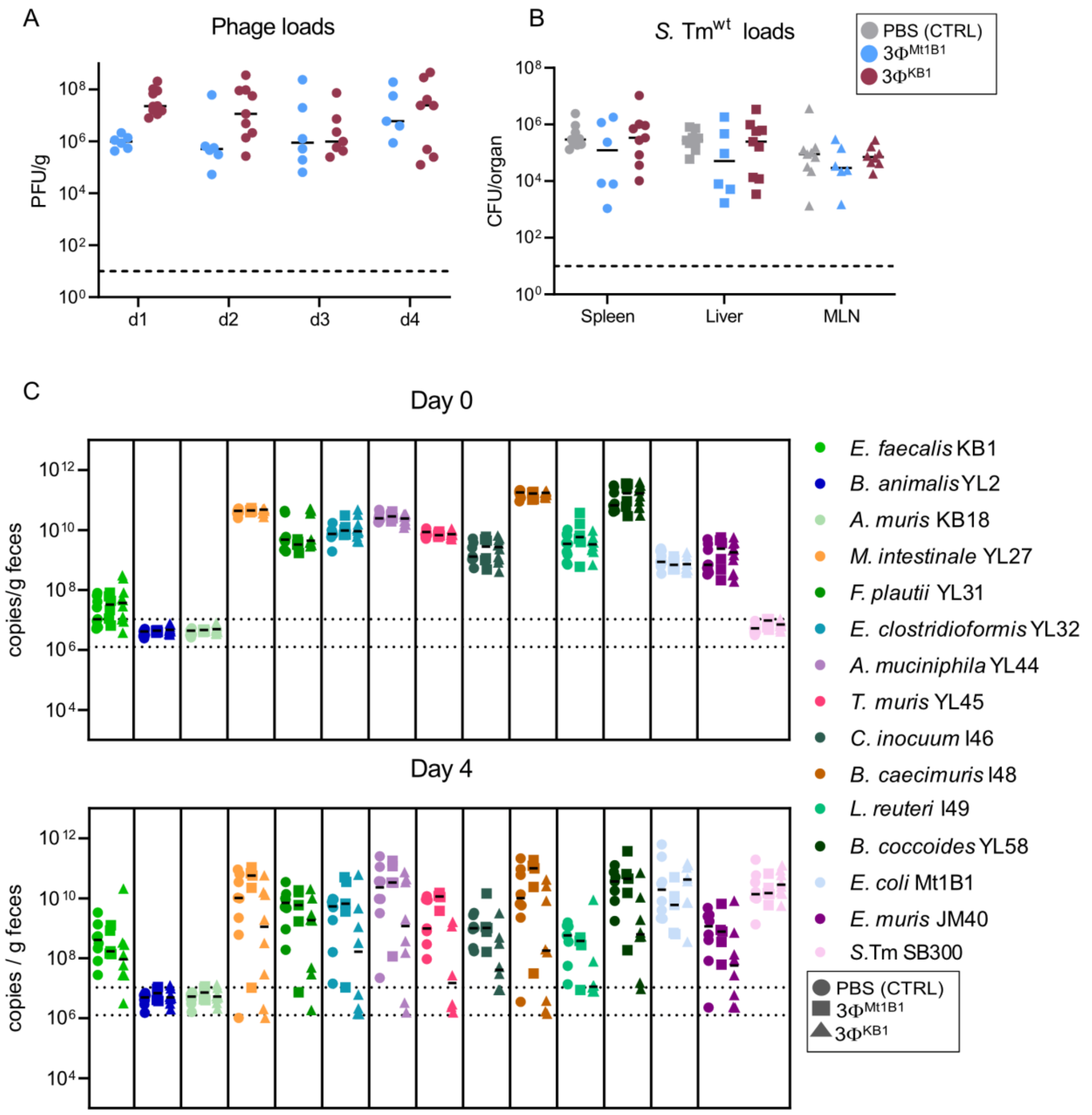


**Fig S6: Microbiota composition after phage challenge in mice infected with *S.* Tm$^{wt}$**

**(A)** Phage loads in feces from experiment shown in Fig. 4, determined by spot assays. **(B)** *S.* Tm$^{wt}$ loads in spleen, liver and mesenteric lymph nodes (MLN), determined by plating. (C) absolute abundance of all 14 bacteria on day 0 and day 4 p. i. with S. Tmwt, determined by strain-specific qPCR. Each dot represents one mouse, black line indicates median, dotted lines indicate DTL.

Table S1

| Designation | Sequence (5´- 3´) | Specificity | Reference |
|---|---|---|---|
| vB_efa_Str1_std_fwd | GGCAAGAACTTATGGAACA | vB_efa_Str1 (major capsid protein) | This study |
| vB_efa_Str1_std_rev | CTGGGTGTCAAAGTGATAA | | |
| vB_efa_Str2_std_fwd | AACTTACTGGCAACTGAC | vB_efa_Str2 (major capsid protein) | This study |
| vB_efa_Str1_std_rev | TACCTTTTCTTCTGGCTCT | | |
| vB_efa_Str6_std_fwd | TTTACCTCAATGTCCACC | vB_efa_Str6 (major tail protein) | This study |
| vB_efa_Str6_std_rev | TCTAGCTACATTCGTGGT | | |
| Mt1B1_P3_fwd | GTCTGGCTTCGATTCTTT | Mt1B1_P3 (phage tail fibers protein) | This study |
| Mt1B1_P3_rev | GGCTTTTCTACTTCCCTG | | |
| Mt1B1_P10_fwd | ATCCCACCTCCTTATGCT | Mt1B1_P10 (phage tail fibers protein) | This study |
| Mt1B1_P10_rev | GTACGCAAGTAACCTATCCC | | |
| Mt1B1_P17_fwd | CTCGGTAACGTCCACACTA | Mt1B1_P17 (phage tail fibers protein) | This study |
| Mt1B1_P17_rev | TCGTTGTGGCTTACCTCT | | |
| vB_efa_Str1_fwd_qPCR | AGAAACACGTGCATTACCAGAATC | vB_efa_Str1 (major capsid protein) | This study |
| vB_efa_Str1_rev_qPCR | TCTGGGATAATTGCTGATGCAT | | |
| vB_efa_Str1_Probe | FAM-TTTGAAGGTGTTAAGTCTG-BHQ-1 | | |
| vB_efa_Str2_fwd_qPCR | GCCCTAAACAACTACAACCATGAA | vB_efa_Str2 (major capsid protein) | This study |
| vB_efa_Str2_rev_qPCR | CTGAAGACCAGTTCTCTCCCAAA | | |
| vB_efa_Str2_Probe | HEX-TTGGTGCAGCTTGGA-BHQ1 | | |
| vB_efa_Str6_fwd_qPCR | CGCCTCGTTGTGCTGCTA | vB_efa_Str6 (major tail protein) | This study |
| vB_efa_Str6_rev_qPCR | CGTGGTACGGCAGTATTAATCG | | |
| vB_efa_Str6_Probe | HEX-ATCCATTCGCCAAGGTCGTTCTGTACC-BHQ1 | | |
| P3_fwd_qPCR | GTAATCTGTGCGCCAGTCGTT | Mt1B1_P3 (phage tail fibers protein) | This study |
| P3_rev_qPCR | CAGGGCAGCGCACCAT | | |
| P3_probe | FAM-AACTGGTGCCTTCACCTCCGCAAAA-BHQ1 | | |
| P10_fwd_qPCR | GCGATCGTGTACCAAGGGATA | Mt1B1_P10 (phage tail fibers protein) | This study |
| P10_rev_qPCR | GGATATTGAGATTGCTGGCCTTA | | |
| P10_probe | HEX-TCTGTGCGCAATACCAGAAGTCATACCTGC_BHQ1 | | |
| P17_fwd_qPCR | GCGCAGACATGTGACTTGTAAAG | Mt1B1_P17 (phage tail fibers protein) | This study |
| P17_rev_qPCR | GATAACAACGAAGGAAGAACACCAA | | |
| P17_probe | HEX-CGCAGCCACTTCTCCGTTGGGA_BHQ1 | | |

Table S2

| Plasmid | Backbone | Origin of insert | Restriction site used for linearization | Reference |
|---|---|---|---|---|
| pAvS1 | pJET 1.2 | vB_efa_Str1 | NotI | This study |
| pAvS2 | pJET 1.2 | vB_efa_Str2 | NotI | This study |
| pAvS3 | pJET 1.2 | vB_efa_Str6 | NotI | This study |
| pAvS4 | pJET 1.2 | Mt1B1_P3 | NotI | This study |
| pAvS5 | pJET 1.2 | Mt1B1_P10 | NotI | This study |
| pAvS6 | pJET 1.2 | Mt1B1_P17 | NotI | This study |
